# Protein sequence classification using natural language processing techniques

**DOI:** 10.1101/2024.08.23.609306

**Authors:** Huma Perveen, Julie Weeds

**Affiliations:** School of Mathematical and Physical Sciences, University of Sussex, UK; School of Engineering and Informatics, University of Sussex, UK

**Keywords:** Protein Sequence Classification, BERT Model, Deep Learning, Bioinformatics, Natural Language Processing, Protein Data Bank (PDB), Sequence Tokenization, Model Fine-Tuning, Hyperparameter Optimization, Machine Learning in Proteomics

## Abstract

Proteins are essential to numerous biological functions, with their sequences determining their roles within organisms. Traditional methods for determining protein function are time-consuming and labor-intensive. This study addresses the increasing demand for precise, effective, and automated protein sequence classification methods by employing natural language processing (NLP) techniques on a dataset comprising 75 target protein classes. We explored various machine learning and deep learning models, including K-Nearest Neighbors (KNN), Multinomial Naïve Bayes, Logistic Regression, Multi-Layer Perceptron (MLP), Decision Tree, Random Forest, XGBoost, Voting and Stacking classifiers, Convolutional Neural Network (CNN), Long Short-Term Memory (LSTM), and transformer models (BertForSequenceClassification, DistilBERT, and ProtBert). Experiments were conducted using amino acid ranges of 1-4 grams for machine learning models and different sequence lengths for CNN and LSTM models. The KNN algorithm performed best on tri-gram data with 70.0% accuracy and a macro F1 score of 63.0%. The Voting classifier achieved best performance with 74.0% accuracy and an F1 score of 65.0%, while the Stacking classifier reached 75.0% accuracy and an F1 score of 64.0%. ProtBert demonstrated the highest performance among transformer models, with a accuracy 76.0% and F1 score 61.0% which is same for all three transformer models. Advanced NLP techniques, particularly ensemble methods and transformer models, show great potential in protein classification. Our results demonstrate that ensemble methods, particularly Voting Soft classifiers, achieved superior results, highlighting the importance of sufficient training data and addressing sequence similarity across different classes.

**Author summary:** Proteins are vital to numerous biological functions, with their sequences determining their roles within organisms. Traditional methods of protein function determination are time-consuming. This study proposes a system utilizing machine learning, deep learning, and natural language processing (NLP) techniques for protein sequence classification. We explore various models, including K-Nearest Neighbors, Multinomial Naive Bayes, Logistic Regression, Multilayer Perceptron, Decision Tree Classifier, Random Forest Classifier, XGBoost, ensemble methods (Voting and Stacking classifier), LSTM, CNN, and transformer models (BertForSequenceClassification, DistilBERT, ProtBert). Our results demonstrate that ensemble methods, particularly Voting Soft classifiers, achieve superior performance, highlighting the potential of advanced NLP techniques in protein classification.

## 1. Introduction

Proteins play crucial roles in living organisms, including catalyzing metabolic processes, replicating DNA, reacting to stimuli, providing structure, and transporting molecules [1]. Proteins are composed of long chains of amino acids, and the sequence of these amino acids determines the protein’s structure and function [2]. Understanding the relationship between amino acid sequence and protein function has significant scientific implications, such as identifying errors in biological processes and clarifying protein synthesis mechanisms.

The study of proteins and other molecules to ascertain the function of many novel proteins has become the foundation of contemporary biological information science. Various methods have been developed to encode biological sequences into feature vectors and classify them using machine learning algorithms. In an experiment, Dongardive et al found that biological sequences are encoded into feature vectors using the N-gram algorithm. 717 sequences divided unevenly into seven classes make up the dataset used for the studies. The closest neighbors are determined using the Euclidean distance and the cosine coefficient similarity metrics [4]. In their 2017 study, Li M. et al. classified the protein sequences of GCPRs (G-protein Coupled Receptors). The dataset included 1019 different protein sequences from the GCPR superfamily. These sequences have all been examined in UniProtKB. The data was pre-processed, and the feature selection methods Term Frequency - Inverse Document Frequency (TF-IDF) and N-gram were utilized [5].

In their work, Lee T. and Nguyen T. used the analysis of unprocessed protein sequences to learn dense vector representation. The information was gathered from the 3,17,460 protein sequences and 589 families in the Universal Protein Resource (UniProt) database. Using Global Vectors for Word Representation (GloVe), a distributed representation was made by encoding each sequence as a collection of trigrams that overlapped [6]. According to Vazhayil A. et al., a protein family is a group of proteins that have the same functions and share similar structures at the molecular and sequence levels. Although a sizable number of sequences are known, it is noted that little is known about the functional characteristics of the protein sequences. Swiss-Prot’s Protein Family Database (Pfam), which has 40433 protein sequences from 30 distinct families, was used as the source of the data for this study. Redundancy in the dataset was checked, and it was discovered that there were no redundant protein sequences. To represent discrete letters as vectors of continuous numbers, the text data was first processed using Keras word embedding and N-gram [7]. Islam et al. developed an NLP-based protein classification method that automates the feature generation process using a modified combination of n-grams and skip-grams (m-NGSG). This approach outperformed existing models in cross-validation accuracy across twelve datasets from nine studies, enhancing the efficiency and accessibility of protein classification from primary sequence data [38].

Given the extensive applications in clinical proteomics and protein bioinformatics, protein sequence analysis has been a focus of numerous studies in recent years, including works by Barve [8], Chen [9], Cong [10], Machado [11], Carregari [12], and Liu [13]. This analysis is crucial for characterizing protein sequences and predicting their structures and functions. Comparative analysis of protein sequences has proven more sensitive than direct DNA comparison, leading to the establishment of multiple protein sequence databases, such as PIR, PDB, and UniProt.

Protein sequence classification, as highlighted by Baldi and Brunak, plays a pivotal role in these analyses because members of the same protein superfamily are often evolutionarily related and share functional and structural similarities. Correctly classifying a protein sequence into its superfamily streamlines molecular analysis within that group, reducing the need for exhaustive individual protein sequence analysis. Typically, two protein sequences are classified together if their features, identified through sequence alignment algorithms, exhibit high homology [14].

Various alignment algorithms have been developed to determine the class of an unknown protein sequence by comparing it with known sequences, calculating similarities, and using databases such as iPro-Class, SAM, and MEME. However, these comparisons can be time-consuming, especially with large databases or long sequences, making it crucial to develop efficient classification systems.

In recent decades, several methods have been employed for general signal classification based on statistical theory, such as decision trees, support vector machines (SVM), and neural networks (NN). Yang et al. utilized word segmentation for feature extraction and SVM for classification [15], while Caragea et al. employed hashing to reduce the dimensionality of protein sequence feature vectors before classification with SVM [16].

Neural networks have also become popular for protein sequence classification due to their ability to handle the high-dimensional, complex features of protein sequences. For example, Wang et al. proposed a modular radial basis function (RBF) neural network for protein sequences [17,18], while Wang and Huang applied a computationally efficient extreme learning machine (ELM) for single-layer feedforward neural networks (SLFNs) to this task [19-22]. Their results indicated that ELM significantly outperformed traditional gradient-based methods which was developed by Levenberg [23] and Marquardt [24] in both speed and classification rate.

To further enhance classification performance while maintaining reasonable training times, Cao et al. introduced a self-adaptive evolutionary ELM (SaE-ELM), which utilizes differential evolution to optimize the hidden neuron parameters in ELM networks [25].

Cao and Xiong investigated the protein sequence classification problem using SLFNs on the protein sequence datasets from the Protein Information Resource center (PIR) and introduced ensemble-based methods to enhance classification performance. Their study proposed two novel algorithms, voting based ELM (V-ELM) and voting based optimal pruned ELM (VOP-ELM), which outperformed existing state-of-the-art methods in accuracy. Particularly, VOP-ELM demonstrated the highest recognition rate, making it a significant advancement in protein sequence classification [26].

A few recent studies have pretrained deep neural language models on protein sequences e.g. Evolutionary Scale Modeling (ESM) [27], Tasks Assessing Protein Embeddings (TAPE-Transformer) [28], ProtTrans [29]. By training both auto-regressive and auto-encoder models on massive protein datasets, recent studies have demonstrated that these protein-specific LMs (pLMs) can capture biophysical properties and make highly accurate predictions in tasks such as protein secondary structure and sub-cellular location, often outperforming traditional methods. This progress highlights the potential of pLMs in advancing protein sequence analysis [29].

Shinde et al considered the structural protein sequences with 10 classes and were able to get 90 % accuracy using a convolutional neural network [30]. A ResNet-based protein neural network architecture is proposed by Maxwell et al. for Pfam dataset families [31]. ProteinBERT is a deep language model created exclusively for proteins, according to Nadav et al. ProteinBERT’s architecture combines local and global representations, enabling the processing of inputs and outputs from beginning to end. Even with limited labelled data, ProteinBERT offers an effective framework for quickly developing protein predictors [32]. The ProtBert model was pre-trained on Uniref100 [33] a dataset consisting of 217 million protein sequences [34].

Traditional methods for determining protein functions, such as crystallography and biochemical studies, are time-consuming [36]. To enhance protein classification, we propose a system that utilizes machine learning, deep learning, and NLP techniques. NLP has emerged as a powerful tool for classifying protein sequences. By treating protein sequences like text, NLP techniques, such as n-grams and word embeddings, enable efficient classification and functional prediction. Models like BERT enhance these processes by learning contextual relationships within the sequences, significantly improving the extraction of valuable insights from complex biological data.

## 2. Material and methods

### 2.1 Dataset

The dataset used in this study is the structural protein sequences dataset from Kaggle [35], derived from the Protein Data Bank (PDB) of the Research Collaboratory for Structural Bioinformatics (RCSB). It contains over 400,000 protein structural sequences, organized into two files: ‘pdb_data_no_dups.csv‘ (protein metadata) and ‘data_seq.csv‘ (protein sequences).

### 2.2 Exploratory data analysis (EDA)

The initial preprocessing steps involved merging the two CSV files on the structure ID column, removing duplicates, dropping null values, and selecting data where the macromolecule type is protein.

#### 2.2.1 Sequence length analysis

After analyzing the statistics in Fig 1a and observing the boxplot in Fig 1b, it can be concluded that the mean sequence character count is approximately 280. This length is considered optimal for applying BERT models, as it provides a sufficient amount of information for deep learning algorithms to process effectively. For the purpose of this study, sequences with more than 30 amino acids in length were considered. This threshold ensures that the sequences are long enough to capture meaningful patterns and interactions. The sequence character count is a crucial parameter in deciding the appropriate sequence length for deep learning applications, impacting the performance and accuracy of the models employed.

**Fig 1.**
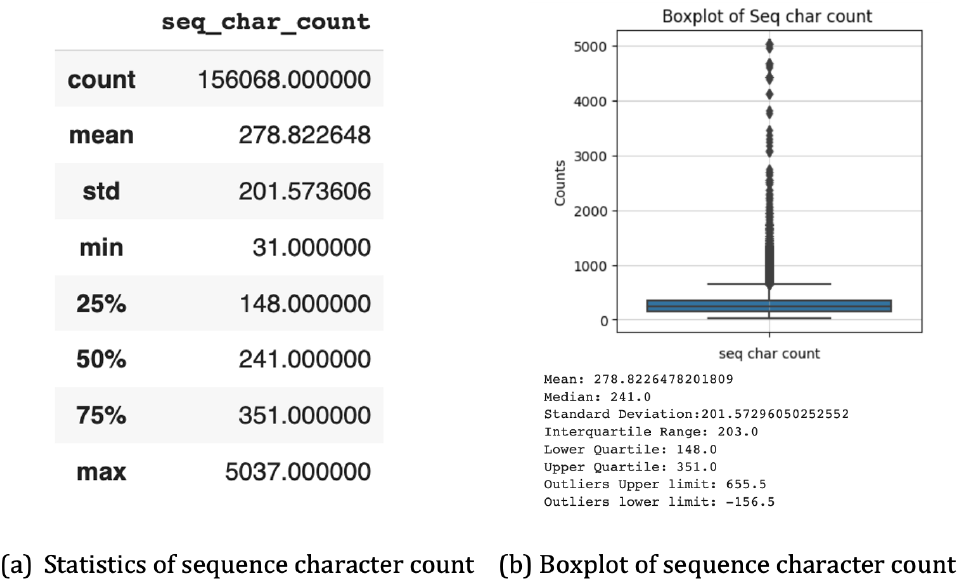
Sequence character count.

In our analysis of the sequence lengths, as depicted in Fig 2, it is evident that the distribution is highly skewed. The majority of unaligned amino acid sequences fall within a character count range of 50 to 450. This skewness indicates a significant variation in sequence lengths within the dataset, which could have implications for downstream analyses and the computational approaches employed. Understanding the distribution of sequence lengths is crucial for optimizing alignment algorithms and improving the accuracy of subsequent bioinformatics analyses.

**Fig 2.**
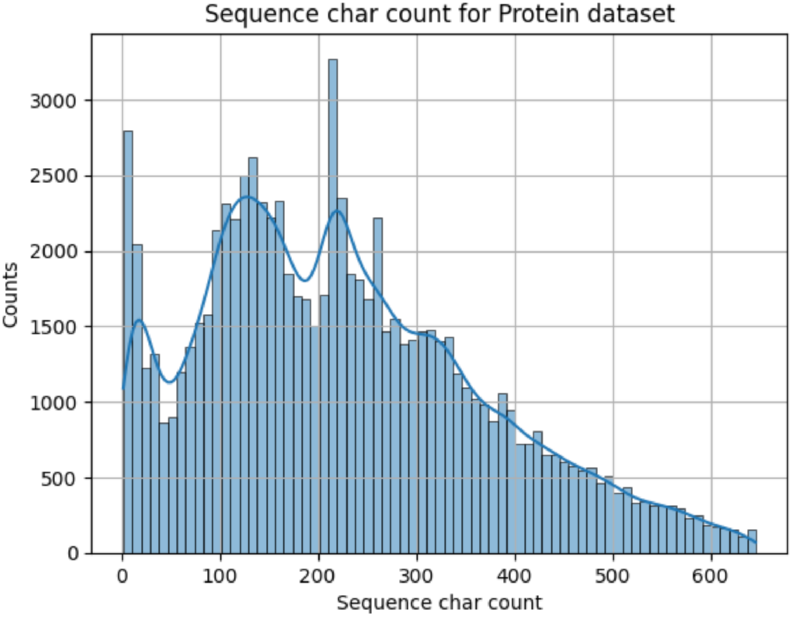
Distribution of sequence length.

#### 2.2.2 Amino acid frequency analysis

Amino acid sequences are represented with their corresponding 1-letter code, for example, the code for alanine is (A), arginine is (R), and so on. The complete list of amino acids with their code can be found in reference [3].

In our analysis, Fig 3 highlights the frequency distribution of amino acids in the dataset. It is evident that leucine (L) appears most frequently, succeeded by alanine (A), glycine (G), and valine (V). This observation aligns with the known biological abundance of these amino acids in various proteins. Additionally, our sequence encoding approach focused on the 20 standard amino acids, deliberately excluding the rare amino acids such as X (any amino acid), U (selenocysteine), B (asparagine or aspartic acid), O (pyrrolysine), and Z (glutamine or glutamic acid) to streamline the analysis and ensure consistency.

**Fig 3.**
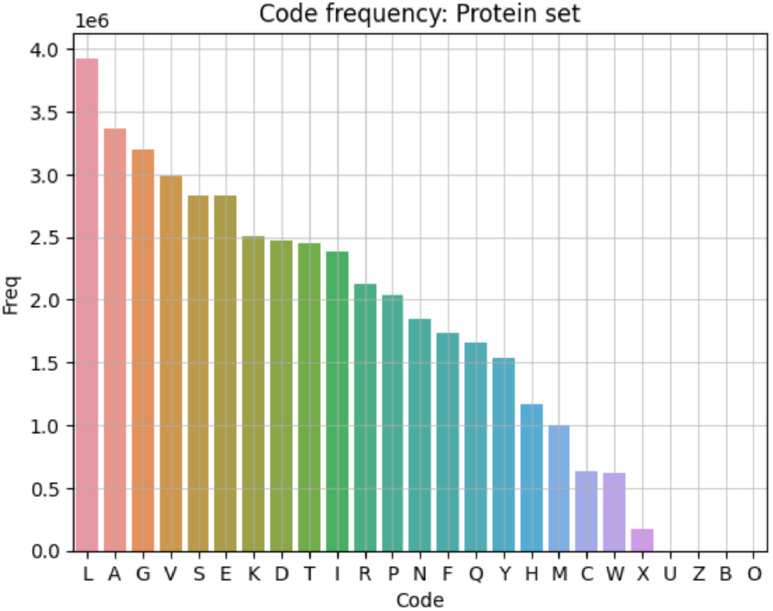
Amino acid distribution.

### 2.3 Machine learning methods

#### 2.3.1 Data transformation

Sequences and classes were initially converted into numerical form using CountVectorizer and LabelEncoder. The dataset was then split into training and test sets with an 80:20 ratio. To optimize the model’s hyperparameters, RandomizedSearchCV was employed, ensuring the best possible configuration for each machine learning method.

#### 2.3.2 Methods

Several machine learning algorithms were applied to the dataset. These included K-Nearest Neighbors (KNN), Multinomial Naive Bayes, Logistic Regression, Multilayer Perceptron, and Decision Tree Classifier. Additionally, various ensemble methods were utilized, such as Random Forest Classifier, XGBoost Classifier, Voting Classifier, and Stacking Classifier, to improve classification performance through model combination and aggregation.

### 2.4 Deep learning methods

#### 2.4.1 Further pre-processing of sequences with Keras

Further pre-processing of sequences was conducted using Keras. The sequences were tokenized, translating each character into a number, and padded to ensure uniform length with maximum lengths of 100, 256, and 512 for evaluating model efficiency and performance. For instance, a sequence ‘GSAFCNLARCELSCRSLGLLGKCIGEECKCVPY’ will be converted into encoding like this [6, 16, 1, 5, 2, 12, 10, 1, 15, 2, 4, 10, 16, 2, 15, 16, 10, 6, 10, 10, 6, 9, 2, 8, 6, 4, 4, 2, 9, 2, 18, 13, 20].

The padded sequence would be like the below:

[6, 16, 1, 5, 2, 12, 10, 1, 15, 2, 4, 10, 16, 2, 15, 16, 10, 6, 10, 10, 6, 9, 2, 8, 6, 4, 4, 2, 9, 2, 18, 13, 20, 0, 0, 0, 0, 0, 0, 0, 0, 0, 0, 0, 0, 0, 0, 0, 0, 0, 0, 0, 0, 0, 0, 0, 0, 0, 0, 0, 0, 0, 0, 0, 0, 0, 0, 0, 0, 0, 0, 0, 0, 0, 0, 0, 0, 0, 0, 0, 0, 0, 0, 0, 0, 0, 0, 0, 0, 0, 0, 0, 0, 0, 0, 0, 0, 0, 0, 0]

The data was split into training, validation, and test sets in a 70:10:20 ratio. For class transformation, sequences with counts greater than 100 were selected, and labels were transformed to one-hot representation using LabelBinarizer. Class weights were assigned using sklearn’s compute_class_weight module to address class imbalance. Early stopping was implemented as a regularization technique to prevent overfitting, with performance monitored after each epoch. The deep learning models used included Long Short-Term Memory (LSTM) and Convolutional Neural Network 1D (CNN 1D).

#### 2.4.2 Long short-term memory (LSTM)

An embedding layer for the mapping input layer is used. Then, a layer of CuDNNLSTM is used for faster implementation. The dropout layer with 20% neuron dropout applied that adds representational capacity to the model and prevents overfitting. Eventually, a dense layer is used as the output layer with softmax activation. The model is trained using categorical cross entropy and is compiled using Adam optimizer. An outline of proposed implementation steps is drawn in Fig 4.

**Fig 4.**
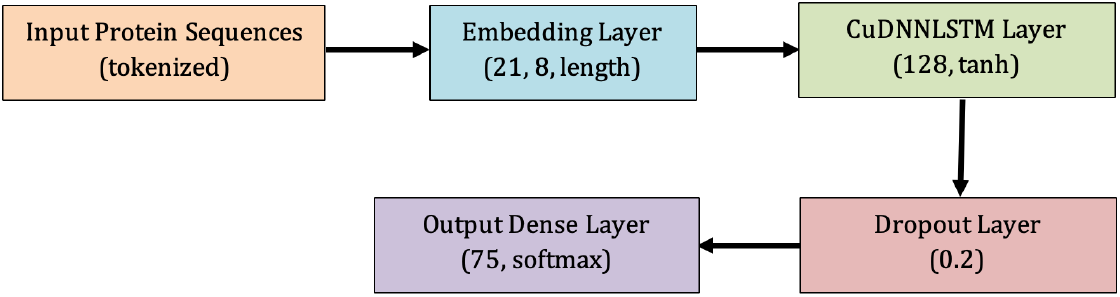
LSTM implementation outline.

#### 2.4.3 Convolutional neural network 1D (CNN 1D)

Recent success in NLP suggests using word embeddings which are already implemented as a Keras Embedding layer. Note that in this dataset, there are only 20 different words (for each amino acid). Instead of using every n-gram, using 1D-convolution on the embedded sequences is considered. The size of the convolutional kernel can be seen as the size of n-grams and the number of filters as the number of words as shown in Fig 5 below.

**Fig 5.**
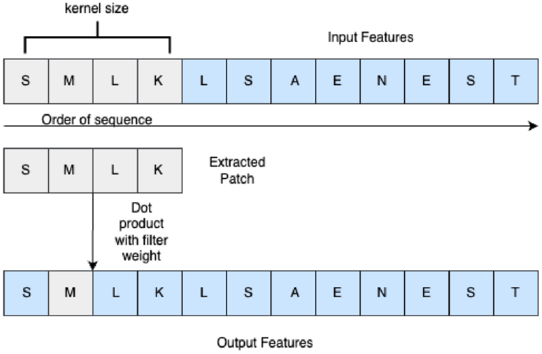
CNN1D mechanism.

A multichannel architecture is used for Convolutional neural network implementation. There are 3 channels. Filter size is 128 and kernel size is 12 in first channel. In second channel filter size is 64 and kernel size is 6 while in third channel filter size and kernel size are 32 and 3 respectively. These filters slide across the input sequence, performing element-wise multiplications and summations to produce feature maps. The filters capture local patterns and dependencies within the sequence of kernel size like n-gram. Each filter learns different features, allowing the model to capture diverse aspects of the input sequence. The filters with different widths help to capture patterns of varying lengths. An activation function ReLU (Rectified Linear Unit) is applied element-wise to introduce non-linearity into the model which sets negative values to zero.

Dropout value is set 0.2, dropout works by “dropping out” a fraction of the neurons in a layer with a specified probability of 0.2. This means that the output of those neurons is temporarily ignored or set to zero. Maxpooling1D layer pool size is kept 2. It selects the maximum value within a fixed window size, reducing the dimensionality of the feature maps while preserving the most salient information. The output of the pooling layers is flattened into a one-dimensional vector. After flattening three channels are concatenated together. Then, a fully connected dense layer is applied that adds representational capacity to the model. Finally, a dense layer is used as the output layer with softmax activation. The model is trained using categorical cross entropy and is compiled using Adam optimizer. An outline of proposed implementation of convolutional neural network is drawn in Fig 6 below.

**Fig 6.**
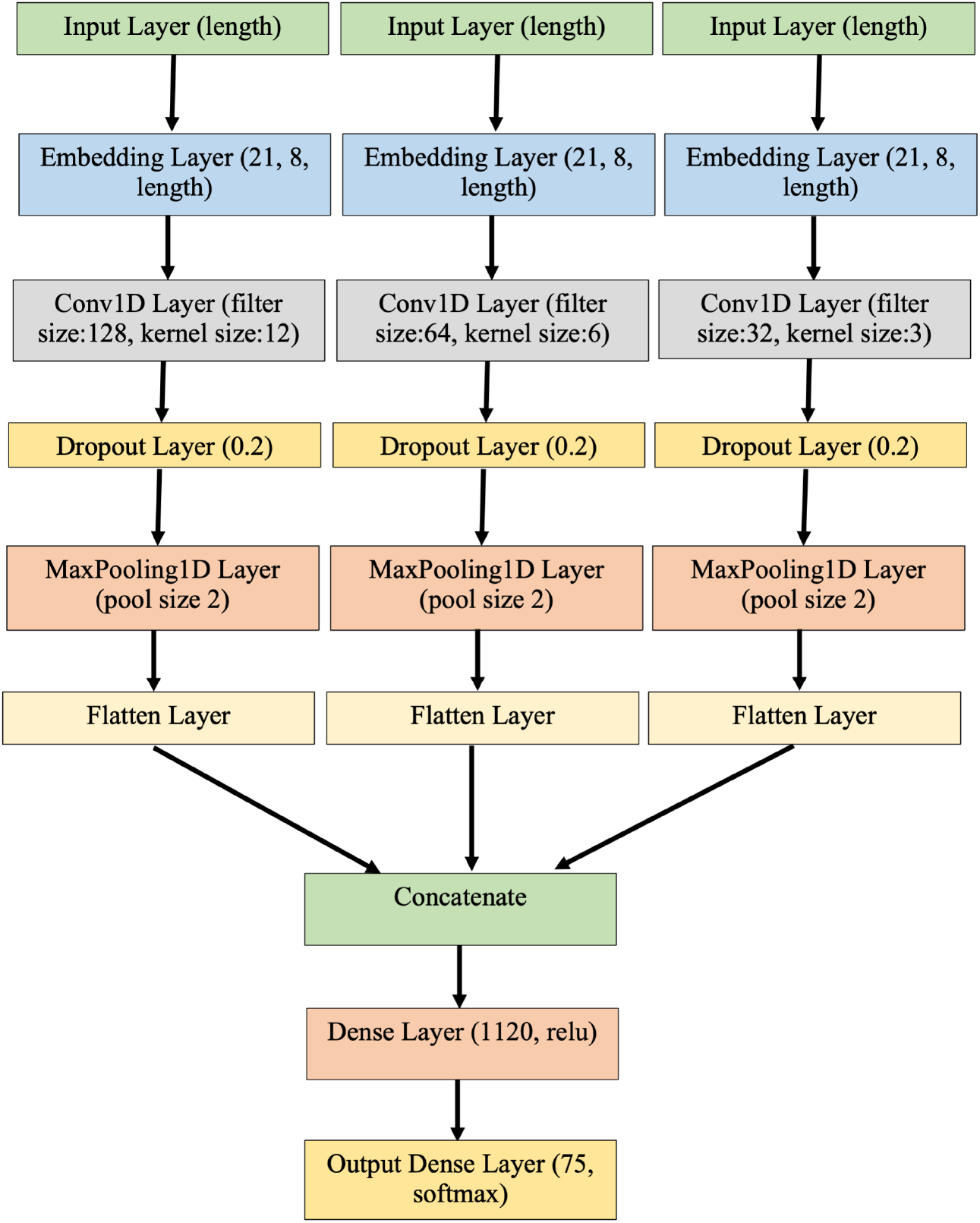
Multi-channel CNN implementation outline.

### 2.5 Transformer models

Three different BERT models were used in this study: BertForSequenceClassification, DistilBERT, and ProtBert.

#### 2.5.1 BertForSequenceClassification

##### Implementation steps

a. **Data pre-processing:** The preprocessed PDB protein dataset, consisting of protein sequences and their corresponding labels, was loaded for analysis. The dataset was split into training, validation, and testing sets (80:10:10) to evaluate the model’s performance. To prepare the protein sequences for input into the model, the BERT tokenizer was used. This tokenizer converts each sequence into a list of tokens, smaller units that BERT can process. The tokens were then converted into input features, including token IDs, attention masks, and segment IDs, which help BERT understand the relationships within the protein sequences.
b. **Model initialization:** The model was initialized with pre-trained weights, capturing knowledge from a large corpus of text data to enhance performance on the protein classification task.
c. **Training loop:** Hyperparameters for training, such as learning rate (2e-5), batch size (4), and number of epochs (30), were set. During each epoch, the model learned from the protein sequences and their corresponding labels in batches. For each batch, a forward pass was performed, obtaining predicted logits. The cross-entropy loss between the predicted logits and true labels was computed to measure performance. A backward pass computed gradients of the loss with respect to the model’s parameters. The optimizer, AdamW, updated the model’s parameters based on the gradients to minimize the loss.
d. **Validation:** After training, the model’s performance was evaluated on the validation set containing unseen protein sequences. A forward pass was performed to obtain predicted logits, which were compared to true labels to assess accuracy, precision, recall, and F1-score.
e. **Fine-tuning and optimization:** If the model’s performance was unsatisfactory, hyperparameters were fine-tuned or different optimization techniques were experimented with. The training loop was repeated with updated settings until the desired performance was achieved.
f. **Evaluation and save model:** Once trained and optimized, the model’s weights and architecture were saved for future use. Evaluation was performed on testing data to determine the model’s performance on unseen data.

#### 2.5.2 DistilBERT

##### Implementation steps

a. **Data preparation and neural network for fine-tuning:** The DistilBert tokenizer fast was used to prepare the protein sequences. A neural network using the DistilBERTClass was created, consisting of the DistilBERT language model, a dropout layer, and a linear layer to produce the final outputs. The protein sequence data was fed into the DistilBERT language model, and the final layer outputs were compared to the encoded categories to determine model accuracy. An instance of this network, referred to as the model, was created for training and subsequent inference.
b. **Loss Function and optimizer:** The cross-entropy loss was calculated to assess performance. An optimizer, AdamW, updated the weights of the neural network to enhance accuracy and efficiency.
c. **c. Model training:** The model was trained over 30 epochs, with a dataloader passing data to the model based on batch size. The output from the model was compared to the actual category to calculate the loss, which was then used to optimize the network’s weights.
d. **Model validation:** During validation, unseen validation data was passed to the model to assess performance and generalization to new data.
e. **Model evaluation:** The model was evaluated using a separate testing dataset to determine performance on entirely new and unseen data, crucial for understanding its real-world applicability.

#### 2.5.3 ProtBert

The implementation steps for ProtBert is same like BertForSequenceClassification.

Below flowchart in Fig 7 shows the experimental setup for three BERT approaches (BertForSequenceClassification, DistilBert and ProtBert).

**Fig 7.**
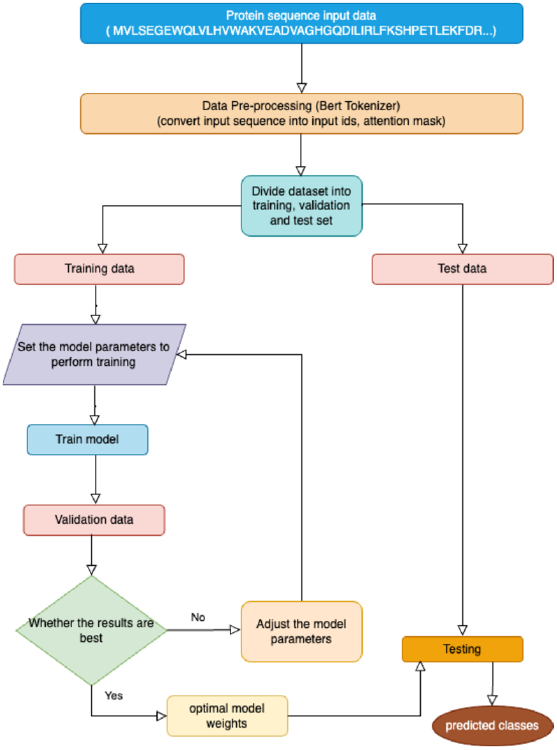
Flow chart for BERT experimental setup.

Parameters used for training Bert models are mentioned in Table 1. All three models were trained over 30 epochs, with performance metrics recorded after every 10 epochs. Due to computational constraints, the training was conducted in three separate intervals. The mean and standard deviation of the accuracy, f1 score and loss over these intervals provide a comprehensive overview of the model’s performance.

**Table 1.**
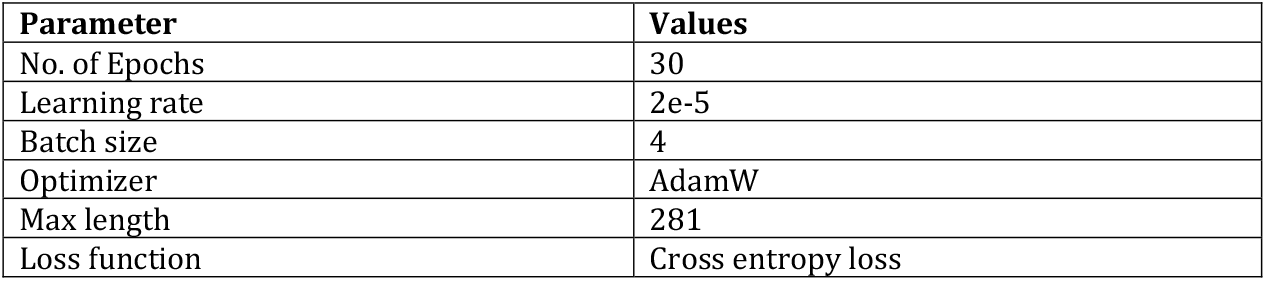
Parameters for Bert models.

| Parameter | Values |
| --- | --- |
| No. of Epochs | 30 |
| Learning rate | 2e-5 |
| Batch size | 4 |
| Optimizer | AdamW |
| Max length | 281 |
| Loss function | Cross entropy loss |

## 3. Results

### 3.1 Machine learning

Training results of best model included in Table 2. The training results of various models demonstrate diverse performance metrics. The **4-gram Voting Soft** model achieved the highest accuracy (0.726 ± 0.006) and weighted F1 score (0.728 ± 0.003), indicating robust overall performance. The **4-gram Stacking** model also performed well, though slightly below Voting Soft in accuracy (0.716 ± 0.006) and weighted F1 score (0.718 ± 0.005). The **2-gram XGB** and **4-gram MLP** models showed strong performance, particularly in accuracy and weighted F1 scores. Conversely, models like **4-gram NB** and **4-gram DTC** exhibited lower performance metrics. Overall, ensemble methods like Voting Soft and Stacking outperformed individual models, showcasing their effectiveness in improving predictive performance across different metrics for imbalanced dataset.

**Table 2.** Result of machine learning approach training.

| Models | Train Accuracy (mean±std) | Train f1-score Weighted (mean±std) | Train f1-score Macro (mean±std) |
| --- | --- | --- | --- |
| **3-gram KNN** | $0.652 \pm 0.003$ | $0.687 \pm 0.018$ | $0.583 \pm 0.002$ |
| **4-gram Multinomial NB** | $0.617 \pm 0.002$ | $0.635 \pm 0.001$ | $0.525 \pm 0.001$ |
| **4-gram LR** | $0.675 \pm 0.003$ | $0.684 \pm 0.003$ | $0.582 \pm 0.002$ |
| **4-gram MLP** | $0.691 \pm 0.001$ | $0.685 \pm 0.001$ | $0.576 \pm 0.006$ |
| **4-gram DTC** | $0.621 \pm 0.001$ | $0.634 \pm 0.002$ | $0.536 \pm 0.004$ |
| **4-gram RFC** | $0.648 \pm 0.001$ | $0.648 \pm 0.001$ | $0.617 \pm 0.002$ |
| **2-gram XGB** | $0.692 \pm 0.002$ | $0.685 \pm 0.002$ | $0.591 \pm 0.002$ |
| **4-gram Voting Soft** | $0.726 \pm 0.006$ | $0.728 \pm 0.003$ | $0.630 \pm 0.009$ |
| **4-gram Stacking** | 0.716 ± 0.006 | 0.718 ± 0.005 | 0.620 ± 0.009 |

Table 3 presents a detailed comparison of different machine learning models’ performance on the test data, analysed across different n-gram ranges (uni-gram, bi-gram, tri-gram, and 4-gram). The evaluation metrics include accuracy, macro F1 score, and weighted F1 score, providing insights into how each model performs under varying textual representations.

**Table 3.** Result of machine learning approach (Test data)

| Metrics | KNN | MultinomialNB | LR | MLP | DTC | RFC | XGB | Voting Classifier | Stacking Classifier |
| --- | --- | --- | --- | --- | --- | --- | --- | --- | --- |
| Uni-gram Accuracy | 0.72 | 0.17 | 0.10 | 0.57 | 0.65 | 0.71 | 0.71 | 0.73 | 0.73 |
| Uni-gram Weighted F1 | 0.71 | 0.17 | 0.09 | 0.55 | 0.65 | 0.71 | 0.7 | 0.73 | 0.73 |
| Uni-gram Macro F1 | 0.60 | 0.1 | 0.09 | 0.4 | 0.53 | 0.62 | 0.61 | 0.63 | 0.61 |
| Bi-gram Accuracy | 0.73 | 0.25 | 0.32 | 0.65 | 0.65 | 0.71 | 0.73 | 0.74 | 0.75 |
| Bi-gram Weighted F1 | 0.73 | 0.26 | 0.32 | 0.65 | 0.65 | 0.71 | 0.73 | 0.74 | 0.74 |
| Bi-gram Macro F1 | 0.60 | 0.24 | 0.29 | 0.49 | 0.53 | 0.63 | 0.64 | 0.64 | 0.62 |
| Tri-gram Accuracy | 0.70 | 0.46 | 0.66 | 0.71 | 0.66 | 0.71 | 0.73 | 0.73 | 0.74 |
| Tri-gram Weighted F1 | 0.74 | 0.48 | 0.67 | 0.71 | 0.66 | 0.71 | 0.72 | 0.73 | 0.74 |
| Tri-gram Macro F1 | 0.63 | 0.47 | 0.59 | 0.6 | 0.55 | 0.63 | 0.64 | 0.64 | 0.62 |
| 4-gram Accuracy | 0.69 | 0.62 | 0.71 | 0.73 | 0.66 | 0.71 | 0.69 | 0.74 | 0.75 |
| 4-gram Weighted F1 | 0.74 | 0.65 | 0.72 | 0.72 | 0.67 | 0.72 | 0.69 | 0.74 | 0.74 |
| 4-gram Macro F1 | 0.63 | 0.55 | 0.62 | 0.6 | 0.55 | 0.63 | 0.61 | 0.65 | 0.64 |

Fig 8 presents a visual comparison of the F1 scores achieved by various machine learning models across different n-gram ranges (uni-gram, bi-gram, tri-gram, and 4-gram). The analysis indicates that models such as K-Nearest Neighbors (KNN), Random Forest, XGBoost, Voting, and Stacking Classifiers maintain consistent F1 scores between 60.0% and 65.0% regardless of the n-gram range, demonstrating their robustness to changes in text representation. The Multi-Layer Perceptron (MLP) shows the lowest performance with the uni-gram model, but its performance improves with higher n-gram ranges, plateauing between the 3-gram and 4-gram models. On the other hand, Multinomial Naïve Bayes and Logistic Regression models exhibit significant improvements in accuracy and F1 score when moving from bi-gram to 3-gram, indicating a strong dependency on n-gram range. Notably, ensemble methods, particularly the Voting Soft classifier, outperform individual models, achieving the highest accuracy (74%), weighted F1 score (74%) and macro f1 score (65%), emphasizing their effectiveness in handling imbalanced datasets.

**Fig 8.**
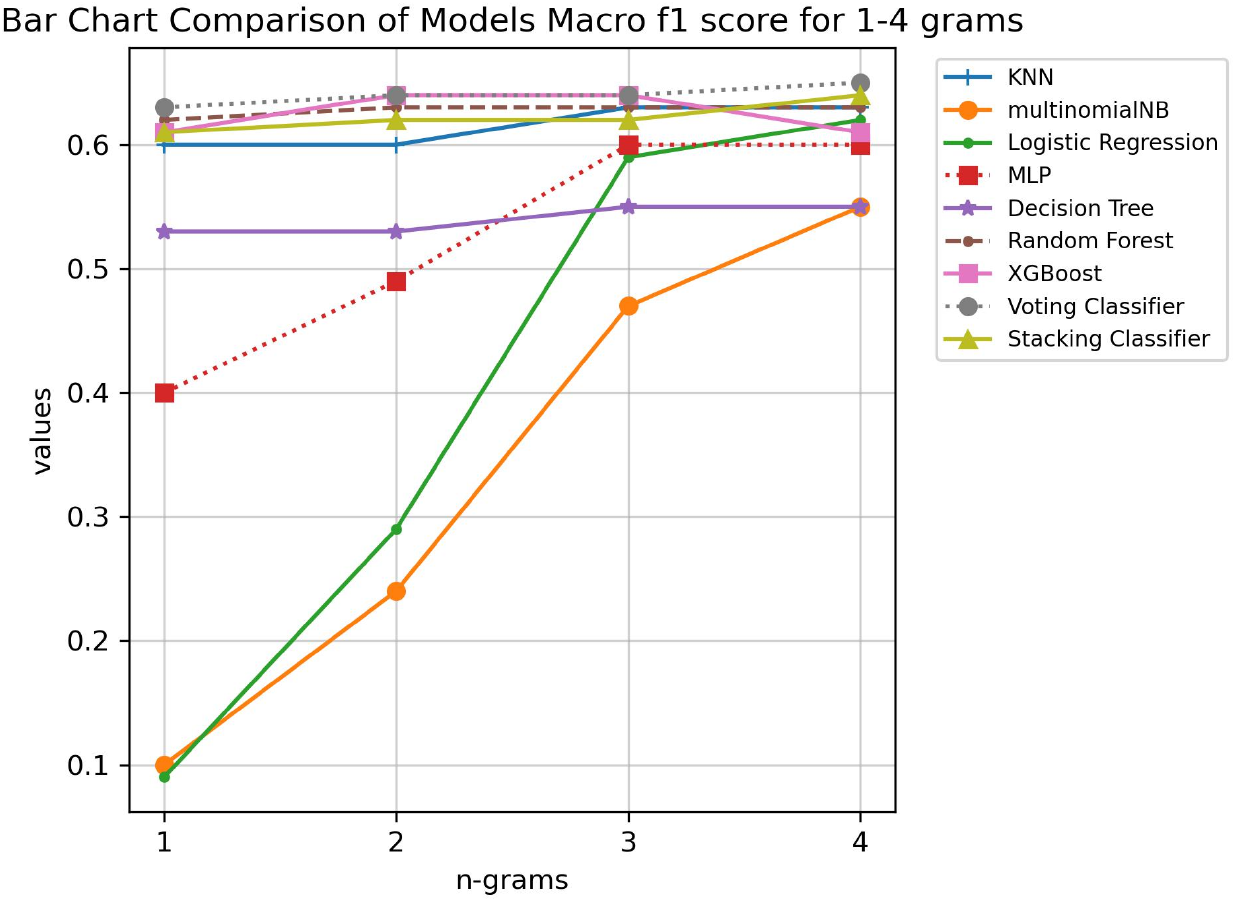
Comparison of models’ macro f1 score for 1-4 grams.

### 3.2 Deep learning

6 experiments are carried out for three different protein sequence lengths with and without class weight for CNN and 3 experiments for LSTM. The LSTM model shows moderate training and validation accuracy, indicating a reasonable but not optimal fit to both the training and validation data. The loss values indicate a moderate level of error in predictions. The F1 scores, particularly the macro F1 score, suggest that while the model performs reasonably well on more frequent classes, its performance on less frequent classes is less reliable and quite variable (Table 4 and Fig 9).

**Table 4.**
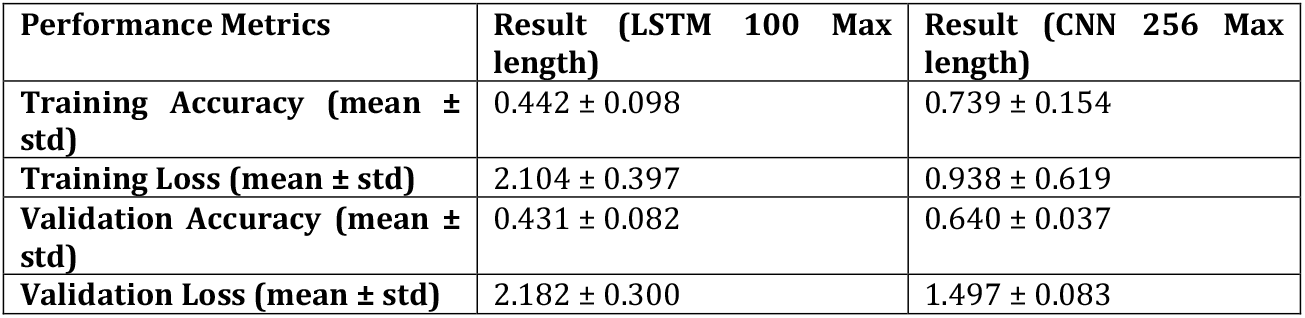
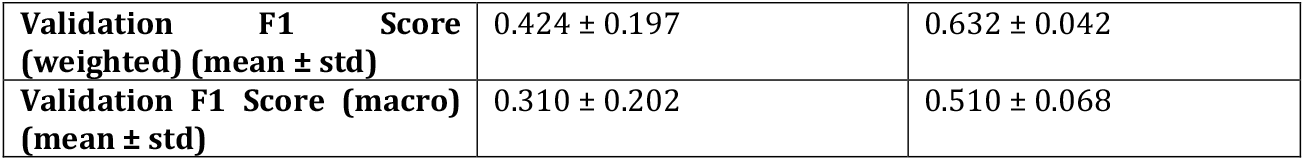
Training result of LSTM and CNN models.

| Performance Metrics | Result (LSTM 100 Max length) | Result (CNN 256 Max length) |
| --- | --- | --- |
| Training Accuracy (mean ± std) | 0.442 ± 0.098 | 0.739 ± 0.154 |
| Training Loss (mean ± std) | 2.104 ± 0.397 | 0.938 ± 0.619 |
| Validation Accuracy (mean ± std) | 0.431 ± 0.082 | 0.640 ± 0.037 |
| Validation Loss (mean ± std) | 2.182 ± 0.300 | 1.497 ± 0.083 |
| **Validation F1 Score (weighted) (mean ± std)** | $0.424 \pm 0.197$ | $0.632 \pm 0.042$ |
| **Validation F1 Score (macro) (mean ± std)** | $0.310 \pm 0.202$ | $0.510 \pm 0.068$ |

**Fig 9.**
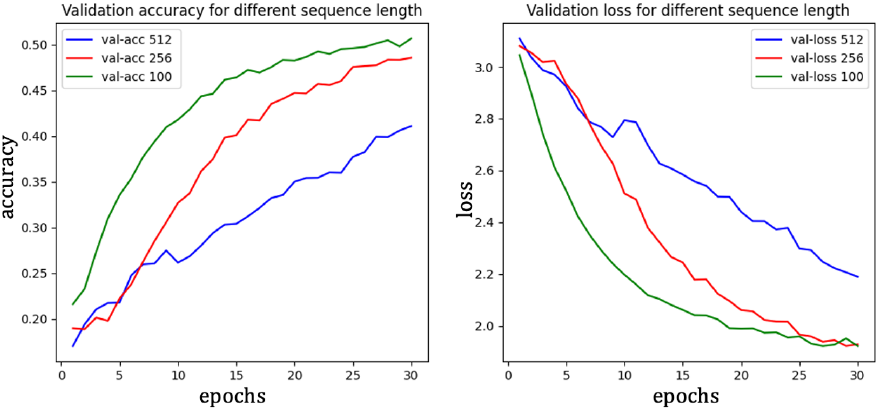
Accuracy and loss for LSTM model for different sequence length (512, 256, 100)

The CNN model shows good training accuracy, indicating effective learning, though with notable performance variability. The training loss reflects accurate predictions but inconsistent errors. Validation metrics indicate moderate generalization with more stable performance than training. F1 scores suggest the model handles frequent classes well, but performance varies across different classes (Table 4 and Fig 10).

**Fig 10.**
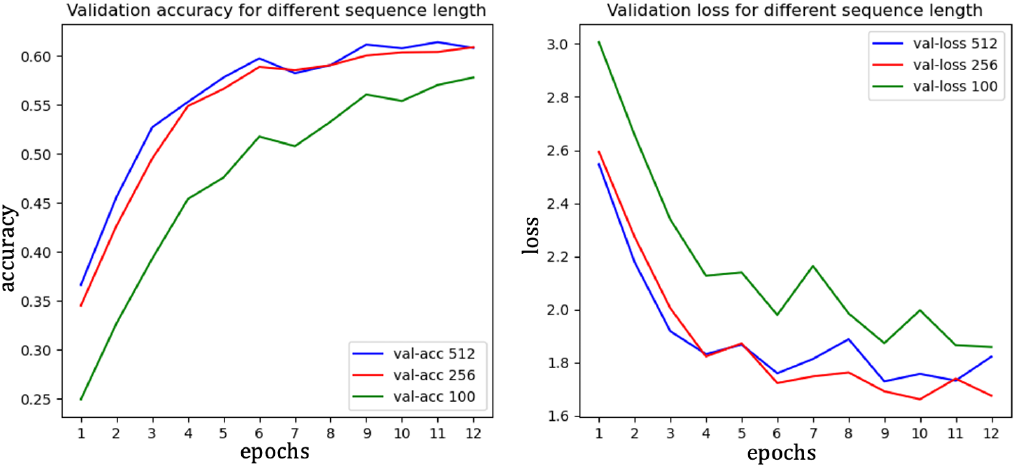
Accuracy and loss for CNN model for different sequence length (512, 256, 100)

Fig 10 illustrates that the CNN model’s accuracy and loss trends are consistent for sequence lengths of 256 and 512, indicating stable model performance across these lengths. However, when the sequence length is reduced to 100, the model struggles to accurately classify protein sequences, resulting in higher loss values. This increased loss suggests a significant drop in model performance, highlighting that shorter sequence lengths are less effective for identifying the correct class in protein sequence classification tasks. The data suggests that longer sequences provide more information, leading to more reliable model predictions and better overall performance.

According to Table 5, The CNN model consistently outperformed the LSTM model across different sequence lengths, with the optimal performance observed at a sequence length of 256. At this length, the CNN achieved an accuracy of 67%, a test loss of 1.48, and a macro F1 score of 55%, making it the best performer among the configurations tested. When class weights were applied, the CNN’s accuracy slightly decreased to 61%, but the macro F1 score improved marginally to 53%. In contrast, the LSTM model showed its best performance at a shorter sequence length of 100, where it achieved an accuracy of 51.0%, a macro F1 score of 48.0%, and a test loss of 1.92. However, the LSTM’s performance deteriorated as the sequence length increased, with accuracy dropping to 41.0% and the macro F1 score falling sharply to 16.0% at a length of 512.

**Table 5.**
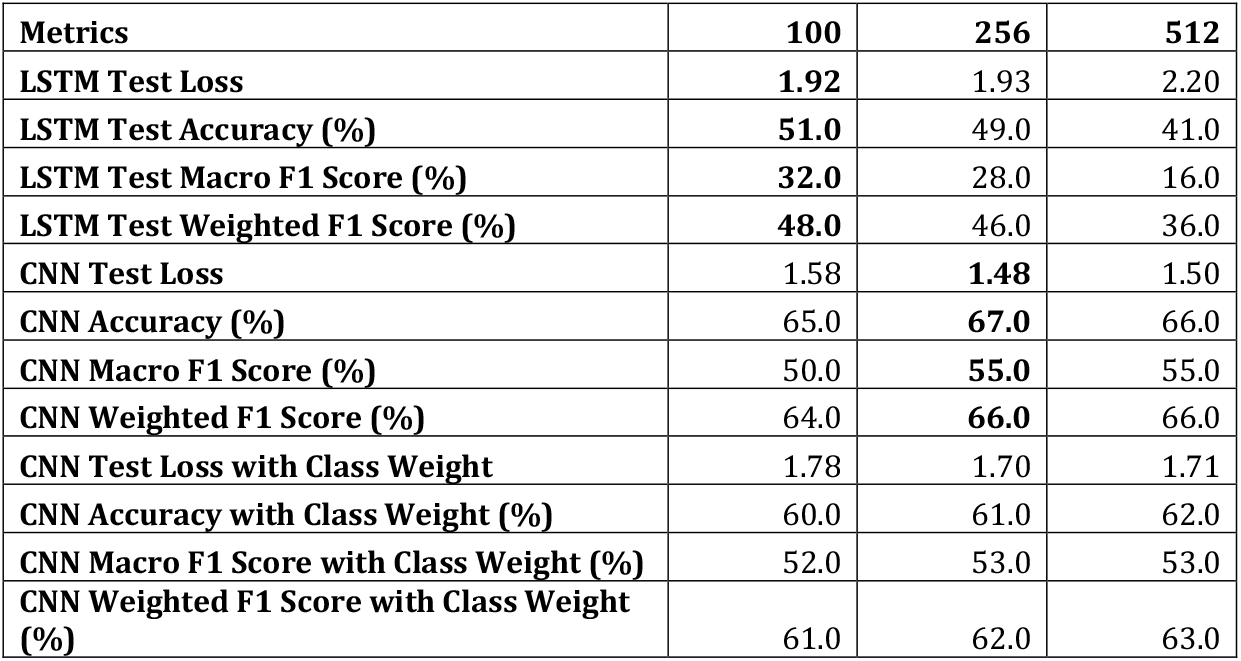
Result of LSTM and CNN for three different sequence length on test data.

| Metrics | 100 | 256 | 512 |
| --- | --- | --- | --- |
| **LSTM Test Loss** | **1.92** | 1.93 | 2.20 |
| **LSTM Test Accuracy (%)** | **51.0** | 49.0 | 41.0 |
| **LSTM Test Macro F1 Score (%)** | **32.0** | 28.0 | 16.0 |
| **LSTM Test Weighted F1 Score (%)** | **48.0** | 46.0 | 36.0 |
| **CNN Test Loss** | 1.58 | **1.48** | 1.50 |
| **CNN Accuracy (%)** | 65.0 | **67.0** | 66.0 |
| **CNN Macro F1 Score (%)** | 50.0 | **55.0** | 55.0 |
| **CNN Weighted F1 Score (%)** | 64.0 | **66.0** | 66.0 |
| **CNN Test Loss with Class Weight** | 1.78 | 1.70 | 1.71 |
| **CNN Accuracy with Class Weight (%)** | 60.0 | 61.0 | 62.0 |
| **CNN Macro F1 Score with Class Weight (%)** | 52.0 | 53.0 | 53.0 |
| **CNN Weighted F1 Score with Class Weight (%)** | 61.0 | 62.0 | 63.0 |

From the Fig 11, it is clearly visible that CNN model without class weight demonstrates superior performance in terms of accuracy and F1 scores, indicating better overall classification capability. The CNN model with class weight shows a slight decrease in performance, suggesting that while class weighting can help in addressing class imbalances, it may also introduce complexities that reduce overall effectiveness. The LSTM model underperforms compared to both CNN models, highlighting its limitations in identifying hidden patterns in longer sequences.

**Fig 11.**
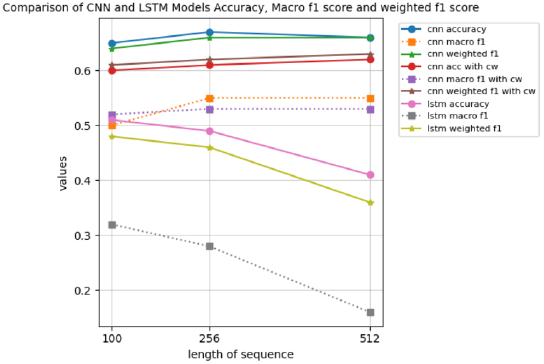
Comparison of deep learning approach.

### 3.3 Transformer models

The training results for BERT models are summarized in Table 6. The Bert For Sequence Classification model demonstrated strong performance during training, with an average loss of 0.3674 and accuracy of 90.73%. However, it exhibited a noticeable drop in validation performance, where the validation loss increased to 1.7626, and accuracy and F1 weighted score dropped to approximately 72%. The DistilBERT model showed a slightly lower training performance, with a loss of 0.4433 and accuracy of 85%. Its validation accuracy was close to Bert For Sequence Classification’s at 72%, with a validation loss of 1.4967, and F1 score of 71.56%. The ProtBert model, while having the highest training loss at 0.7904, demonstrated the best validation performance among the three, with a validation accuracy of 77.4% and an F1 weighted score of 75.42%, indicating its robustness on unseen data. These models achieved a good balance between training and validation performance up to 20 epochs, but beyond that, the improvement in training accuracy did not translate into better validation performance, suggesting the risk of overfitting.

**Table 6.**
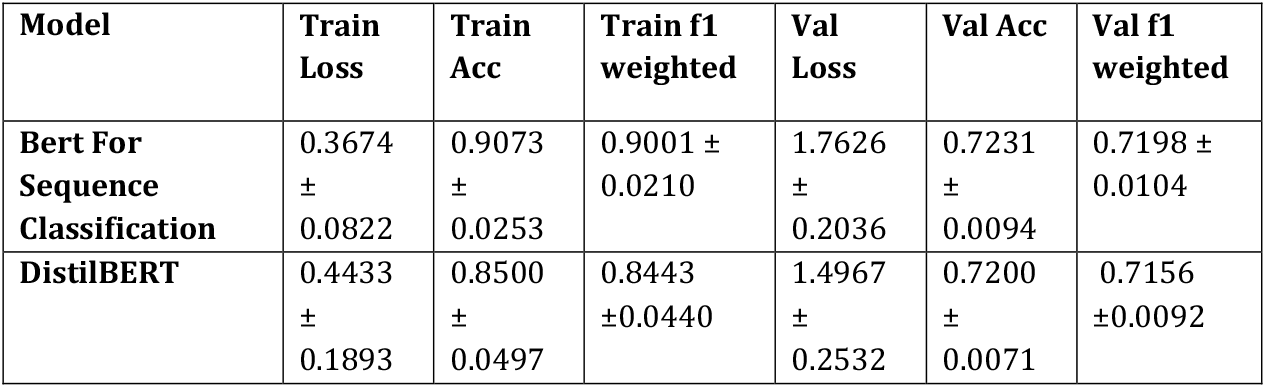
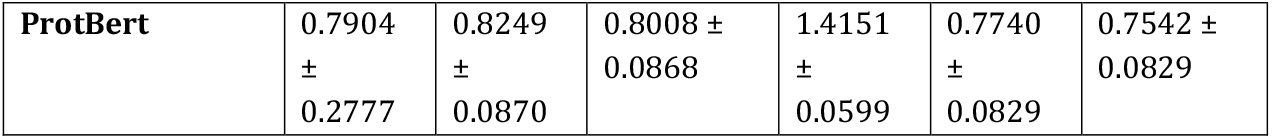
Training result for BERT models.

Table 7 and Fig 12 presents the performance of three different BERT-based transformer models on the test dataset, highlighting key metrics such as test loss, accuracy, and F1 scores. All three models show competitive accuracy, with ProtBert achieving the highest at 77%. BertForSequenceClassification and DistilBERT both have an accuracy of 73%. ProtBert has the lowest test loss (1.37), indicating better performance in terms of prediction error. DistilBERT shows a lower test loss (1.48) compared to BertForSequenceClassification (1.78).

**Table 7.** Test result for BERT models.

| Transformer Model | Test Loss<br>(mean±std) | Accuracy<br>(%)<br>(mean±std) | F1 score<br>weighted<br>(%)<br>(mean±std) | F1 score<br>macro (%)<br>(mean±std) |
| --- | --- | --- | --- | --- |
| **BertForSequenceClassification** | 1.7767±0.2164 | 0.7333±0.0094 | 0.7300±0.0082 | 0.6053±0.0249 |
| **DistilBERT** | 1.4767±0.2368 | 0.7267±0.0125 | 0.7233±0.0094 | 0.6053±0.0309 |
| **ProtBert** | 1.3713±0.1012 | 0.7698±0.0067 | 0.7552±0.0083 | 0.6067±0.0205 |

**Fig 12.**
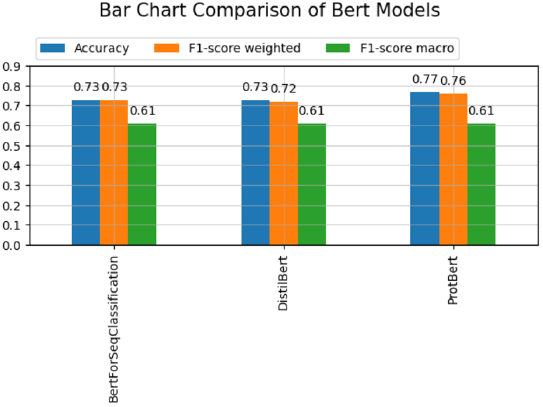
Comparison of Bert models.

ProtBert again leads with a weighted F1 score of 76%, indicating it performs well across classes considering the class distribution. BertForSequenceClassification and DistilBERT had similar performances with slight variations. BertForSequenceClassification and DistilBERT have similar weighted F1 scores, 73% and 72%, respectively. All models have the same macro F1 score of 61%, reflecting their balanced performance across all classes, irrespective of class distribution.

### 3.4 Error analysis

In general, proteins can be different types of enzymes, signalling proteins, structural proteins, and a variety of other options. Since many proteins are designed to bind in the same locations as one another, they frequently exhibit extremely similar properties. A Hydrolase enzyme and a Hydrolase inhibitor protein, for instance, will have similar structures since they focus on the same regions. Fig 13 shows some example of error analysis.

**Fig 13.**
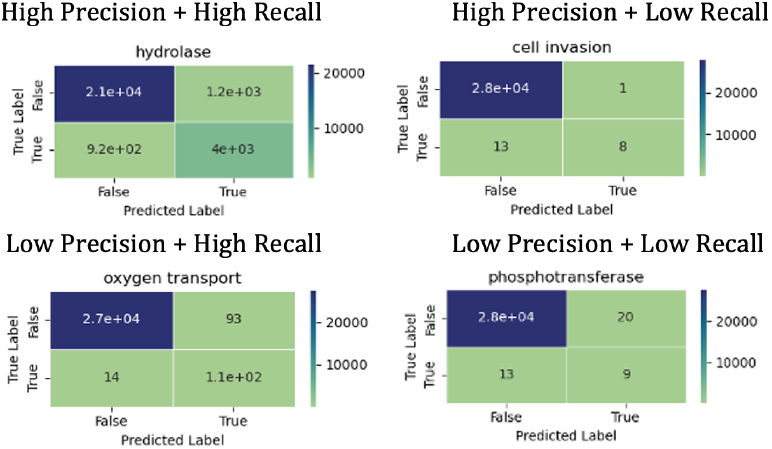
Error analysis.

**high precision and high recall** that means model is able to identify them because these classes either have enough sample or their structure are not similar to others for instance, Allergan, apoptosis, immune system, isomerase, hydrolase.

Classes have **high precision and low recall** that means model is not identify them correctly, example RNA binding proteins, DNA binding proteins, and transcription proteins, all share characteristics with gene regulator proteins and Cell invasion, Cell adhesion having similarity with cell cycle that make model difficult to identify them.

Classes are showing **low precision and high recall** like electron transport and oxygen storage, model is able to well detect these classes but also includes observations from other classes as well, for instance oxygen storage might be make model misleading for oxygen transport.

Classes e.g. Phosphotransferase, Transcription inhibitor having **low precision and low recall** which means model is not able to identify correct classes and model is not doing well on the entire test dataset to find correct classes. On the other side some classes like, Ribosome having low precision and low recall because of similarity with ribosomal protein (fundamental building blocks for ribosome) that means model is not able to detect correct class.

## 4. Discussion

The study demonstrates that NLP techniques can significantly enhance protein sequence classification. The use of n-grams proved effective in improving classifier performance, and ensemble methods showcased their potential in handling imbalanced datasets. While CNN outperformed LSTM in handling longer sequences, transformer models, particularly ProtBert, demonstrated superior accuracy and F1 scores, albeit with higher computational requirements.

The findings of this study highlight the significant potential of natural language processing (NLP) techniques in protein sequence classification. Given the growing amount of biological data, efficient and automated methods are essential. The results demonstrate that various machine learning models, when applied to amino acid n-grams, can achieve noteworthy accuracy and F1 scores, underscoring the effectiveness of this approach.

The K-Nearest Neighbors (KNN) algorithm performed particularly well on tri-gram data, indicating its strength in capturing local sequence similarities. In contrast, models like Logistic Regression, MLP, and Random Forests showed their highest performance with 4-gram data, suggesting that these models benefit from a broader contextual understanding provided by longer n-grams.

The transformer models, particularly ProtBert, showed promise with competitive F1 scores, although they required significant computational resources. This emphasizes the importance of access to high-performance computing facilities for training such models efficiently. The challenges faced due to limited GPU access and session expiry constraints highlight a practical limitation in the current study, suggesting a need for more robust computational infrastructure for future research.

Despite these promising results, several challenges remain. The primary source of error across models was the imbalanced dataset, with some classes having significantly fewer samples. This imbalance likely hindered the models’ ability to generalize well across all classes. Future work could explore advanced techniques for handling class imbalance, such as data augmentation or more sophisticated weighting schemes.

The similarity between sequences from different classes also posed a challenge, potentially confusing the models and reducing classification accuracy. Advanced sequence embedding techniques or incorporating additional biological context could help mitigate this issue.

Furthermore, the results indicate that while CNNs and LSTMs showed reasonable performance, transformer models like BERT variants provided more consistent results across different sequence lengths and configurations. This suggests that transformers may offer a more robust framework for protein sequence classification, benefiting from their ability to capture long-range dependencies and contextual information effectively.

In summary, this study demonstrates the feasibility and effectiveness of using NLP techniques for protein sequence classification. The insights gained here pave the way for further exploration and optimization of these methods, with the potential to significantly enhance our ability to analyze and interpret complex biological data. Future research should focus on addressing the challenges of class imbalance and sequence similarity, as well as leveraging more advanced computational resources to fully realize the potential of these techniques.

## 5. Conclusion

Assigning class weights effectively managed the imbalanced dataset. The best machine learning approach was the Voting Soft classifier, while CNN outperformed LSTM in deep learning methods. Among transformer models, ProtBert achieved the highest performance, highlighting the potential of advanced NLP techniques in protein sequence classification.

As bioinformatics advances uncover novel proteins, the need for precise, effective, and automated protein sequence classification methods becomes increasingly important. Identifying the family or class of a protein sequence, which indicates its function, remains a fundamental challenge in protein sequence analysis. Most previous studies have worked on protein secondary structure and sub-cellular location prediction, which motivated this study to explore various feature extraction methods using natural language processing (NLP) techniques to classify protein sequence.

In this research, we addressed dataset imbalance by assigning class weights. The dataset included 75 target protein classes, and machine learning experiments were conducted using amino acid ranges of 1-4 grams. Among the models tested, the K-Nearest Neighbors (KNN) algorithm excelled with tri-gram data, achieving 70.0% accuracy and a macro F1 score of 63.0%. Multinomial Naïve Bayes achieved 62.0% accuracy and a 55.0% F1 score with 4-gram data. Logistic regression performed best with 4-grams, attaining 71.0% accuracy and a 62.0% F1 score. The Multi-Layer Perceptron (MLP) model reached 73.0% accuracy and a 60.0% F1 score on 4-gram data. The decision tree model achieved 66.0% accuracy and a 55.0% F1 score. The random forest classifier attained 71.0% accuracy and a 63.0% F1 score on 4-gram data. The XGBoost classifier performed well on bi-gram data, with 73.0% accuracy and a 64.0% macro F1 score. The voting classifier achieved 74.0% accuracy and a 65.0% F1 score, while the stacking classifier achieved 75.0% accuracy and a 64.0% macro F1 score.

For convolutional neural network (CNN) and long short-term memory (LSTM) models, experiments were conducted using sequence lengths of 512, 256, and 100. The CNN model showed the best performance with a sequence length of 256, achieving 67.0% accuracy and a 55.0% macro F1 score. The LSTM model had lower accuracy and F1 scores, with the highest scores for a sequence length of 100.

Finally, three transformer models were used: BertForSequenceClassification, DistilBERT, and ProtBert. The results were similar across these models, with BertForSequenceClassification achieving a macro F1 score of 61.0% and a weighted F1 score of 73.0%, DistilBERT achieving 72.0%, and ProtBert achieving 76.0%, although ProtBert required more computational resources. Training ProtBert on a Mac M1 MPS environment took approximately 30 hours for one epoch. Limited GPU access and session expiry issues prevented running the models on Google Colab Pro. Transfer learning on a Mac showed that each epoch for BertForSequenceClassification and DistilBERT required about 5 hours.

The main sources of model error were limited sample training data for some classes and sequence similarity across different classes. Addressing these issues is crucial for further improving protein sequence classification accuracy.

## 6. Acknowledgments

I would like to express my sincere gratitude to my supervisor, Dr. Julie Weeds, for her invaluable guidance, support, and encouragement throughout the course of this research. Her expertise and insights have greatly contributed to the development of this work.

## Data Availability Statements

The dataset is publicly available at https://www.kaggle.com/datasets/shahir/protein-dataset. All source codes used in this study can be found at https://github.com/humaperveen/PortfolioProjects/tree/main/PythonProjects/NLP/ProteinSeqClassification

## Abbreviations

NLP: Natural Language Processing
KNN: K-Nearest Neighbors
Multinomial NB: Multinomial Naive Bayes
LR: Logistic Regression
MLP: Multi-Layer Perceptron
DTC: Decision Tree Classifier
RFC: Random Forest Classifier
XGB: XGBoost Classifier
CNN: Convolutional Neural Network
LSTM: Long Short-Term Memory
PDB: Protein Data Bank
RCSB: Research Collaboratory for Structural Bioinformatics
UniProt: Universal Protein Resource
PIR: Protein Information Resource center
V-ELM: Voting-based Extreme Learning Machine
VOP-ELM: Voting-based Optimal Pruned Extreme Learning Machine
ELM: Efficient Extreme Learning Machine
SLFNs: Single-layer Feedforward Neural Networks
SaE-ELM: Self-adaptive Evolutionary Extreme Learning Machine
ESM: Evolutionary Scale Modeling
TAPE-Transformer: Tasks Assessing Protein Embeddings Transformer
pLMs: Protein-specific Language Models

